# PPI-Rescue: restoration of eye development via functional reconstitution of a transcription complex

**DOI:** 10.64898/2026.11.02.763744

**Authors:** N. Jusić, C. Pina, D. Netz, J. Lee, F. Pignoni, SJ. Neal

## Abstract

The activity of transcription factors (TFs) can be modulated through protein-protein interactions (PPIs) with cofactors which can serve to enhance, repress or alter the specificity of TF function. Current genetic and molecular approaches such as knockout and overexpression studies are likely to affect all possible activities of a TF, presenting a dilemma when attempting to parse out how TFs function within a cell. Hence, there is a need to devise a technique that will allow for mechanistic insight to be gained into the function(s) of discrete transcription complexes. Herein we report “PPI-Rescue”, an approach that leverages bespoke mutations in TFs and the selective reconstitution of specific complexes to study the molecular properties and biological functions of discrete transcription complexes *in vitro* and *in vivo*. We demonstrate PPI-Rescue using the conserved *Drosophila* transcription factor Sine oculis (So) and cofactor Eyes Absent (Eya). To validate this approach, we engineered a missense mutation, SoΔEya (C110S), that selectively abrogates PPI-Eya to show that PPI-Eya is necessary for the transcriptional activity of So in cultured cells and to promote eye formation in *Drosophila*. Using heterologous leucine zipper domains, we specifically reconstituted the SoΔEya-Eya PPI and show that this complex is transcriptionally active in cells and can rescue eye development. PPI-Rescue was also used to restore transcriptional activity to SoΔPPI (V111E), an allele that mimics the human SIX1(V17E) allele associated with Branchio-Oto-Renal syndrome. PPI-Rescue thus represents an innovative approach to understanding the complex molecular logic of transcription complexes in both development and disease.

## Introduction

Transcription factors (TFs) have long been a central focus of molecular genetic studies of development and disease. This has led to the development of numerous techniques to express, knockdown, activate, repress and localize TFs in specific cells and tissues. However, TFs may form discrete complexes with different cofactors in different tissues of an organism, or even within a given cell. These potentially overlapping functions of TFs are difficult to investigate because common molecular genetic techniques typically have global effects on TF and / or cofactor expression and function. In this study, we provide proof-of-concept for an approach to study the role(s) of discrete transcription complexes *in vivo* through the functional reconstitution of specific TF-cofactor molecular interactions. We call this approach “PPI-Rescue”.

A limited repertoire of DNA-binding TFs, numbering ∼1,600 in humans and just ∼750 in the fruit fly *Drosophila* regulate the expression of the genome (Shokri *et al*, 2019). Their roles range from sustaining routine cell functions to guiding the development of complex tissues and organs. The small number of regulators, relative to a much larger number of genes – ∼23,000 genes in humans and ∼14,000 genes in *Drosophila* – necessitates that TFs be repurposed throughout development to fulfill specific roles in the various cell types of the organism (e.g. (Bertrand, 2016; Costa *et al*, 2022; Coux *et al*, 2024; Garcia-Mora *et al*, 2025; Hosokawa *et al*, 2021; Miyares & Lee, 2019; Quan *et al*, 2016)). One mechanism by which TF function can be diversified is through protein-protein interactions (PPIs) with transcription co-activators or co-repressors, collectively “cofactors” or CoFs, which may serve to modulate the specificity and / or the magnitude of TF-dependent transcription (e.g. (Bridoux *et al*, 2020; Carr *et al*, 2015; Lelli *et al*, 2012; Manning *et al*, 2024; Rader *et al*, 2023; Reichlmeir *et al*, 2021)).

The Retinal Determination Gene Network (RDGN) is among the best studied examples of how complex networks of TFs and CoFs can drive the growth and differentiation of a simple epithelium into a complex neural tissue, in this case, the retina (reviewed in (Treisman, 2013)). Genetic analyses of *Drosophila* eye morphogenesis has led to the identification of numerous evolutionarily conserved TFs such as Sine oculis (So) and CoFs such as Eyes Absent (Eya), whose loss of function completely abrogates retinogenesis (Bonini *et al*, 1997; Cheyette *et al*, 1994; Jusiak *et al*, 2014; Serikaku & O’Tousa, 1994; Silver *et al*, 2003; Weasner *et al*, 2007). Sine oculis is the founding member of the SIX family of homeodomain TFs and Eya is the founding member of the namesake EYA family of protein tyrosine phosphatases. In the context of fly retinogenesis, it is postulated that Eya binds to and potentiates the transcriptional activity of So to induce additional RDGN factors (Pignoni *et al*, 1997), including the proneural gene *atonal* (Tanaka-Matakatsu & Du, 2008; Zhang *et al*, 2006), although direct evidence for this physical and functional interaction has not been shown *in vivo*.

In humans, autosomal dominant mutations in SIX1 and EYA1 are pathologically associated with branchio-oto and branchio-oto-renal syndromes, collectively “BOS”, and affect ∼1:40,000 live births (Abdelhak *et al*, 1997; Buller *et al*, 2001; Clark *et al*, 2019; Fraser *et al*, 1980; Kochhar *et al*, 2008; Krug *et al*, 2011; Mehdizadeh *et al*, 2021; Morton & Nance, 2006; Neal *et al*, 2024; Orten *et al*, 2008; Patrick *et al*, 2013; Patrick *et al*, 2009; Ruf *et al*, 2003; Ruf *et al*, 2004; Sanggaard *et al*, 2007; Shah *et al*, 2020; Smith & Azaiez, 1993; Song *et al*, 2013; Vincent *et al*, 1997; Yalcouye *et al*, 2022; Zhang *et al*, 2024a; Zheng *et al*, 2021); SIX1 is an ortholog of *Drosophila* So and EYA1 is an ortholog of *Drosophila* Eya. As the nomenclature suggests, BOS results from the impaired development of multiple organ systems, though it is often initially diagnosed in children that are born congenitally deaf. Informed by the strong history of genetic analyses in the fly, a leading hypothesis for BOS pathogenesis is that mutant protein variants disrupt the assembly or transcriptional activity of the SIX1-EYA1 TF:CoF complex (Krug *et al*., 2011; Patrick *et al*., 2013; Ruf *et al*., 2004).

While the above hypothesis is plausible and is supported by *in vitro* biochemical evidence (Ruf *et al*., 2004), proving that a specific PPI mediates the developmental function of a TF is not so simply demonstrated *in vivo*. For instance, a given TF may have multiple CoFs (e.g. (Ahmed *et al*, 2012; Chen *et al*, 1997; Kenyon *et al*, 2005a; Kenyon *et al*, 2005b; Neilson *et al*, 2010)). In such a case, genetic loss of function studies would necessarily affect the function of all potential complexes comprising that TF. Many putative and confirmed So and SIX1 CoFs have indeed been revealed through unbiased yeast 2-hybrid (Y2H) and affinity-purification mass spectrometry screens (e.g. (Giot *et al*, 2003; Oughtred *et al*, 2021; Stark *et al*, 2006)), suggesting that pleiotropic effects are likely to be observed upon So / SIX1 knockdown or knockout.

To our knowledge, there is currently no reported method by which the function of discrete transcription complexes can be robustly studied *in vivo*. Hence, we set out to develop tools and an *in vivo* model that would allow us to establish the role(s) of a specific transcription complex in development and disease. For this study, we selected the *Drosophila* So-Eya complex as a model. First, we used a Y2H screen and published evidence to identify a series of bespoke So missense variants in which a specific PPI was compromised. Then, we used heterologous leucine zipper domains to selectively reconstitute the disrupted PPI and restore transcription complex function (Moll *et al*, 2001; Pfeiffer *et al*, 2008). Using these approaches, we show herein that So-Eya complex formation is necessary for retinogenesis in *Drosophila* and for transcription activity *in vitro*. We propose that “PPI-Rescue” is a generalizable method to study the potentially nuanced roles of discrete transcriptional complexes in both healthy development and in disease.

## Materials and Methods

### Yeast 2-hybrid (Y2H) analysis

Y2H analysis generally followed the Promega Matchmaker assay protocol. Briefly, Sine oculis bait constructs (So-PA, AA103-276) in pGBKT7 vector were co-transformed with co-factor (Eya, Gro, or Sobp) prey constructs in pActII or pGAD424 vectors into AH109 yeast cells. Co-factor clones used include Eya (Eya-PA, residues 493-766), Gro (Gro-PA, residues 1-247) and Sobp (Sobp-PA, residues 210-749).

Doubly transformed yeast were selected on double dropout media, lacking Trp and Leu. Interactions were tested on triple dropout media (lacking Trp, Leu and His) in the presence or absence of the competitive His biosynthesis inhibitor 3-Amino-1,2,4-triazole (3-AT). Assay stringency was titrated for each bait-prey pair by adjusting the 3-AT concentration to a level (0.1-10 mM) that suppressed the growth of yeast transformed with empty vector and the complementary So bait or CoF prey construct. A positive interaction was scored when yeast co-transformed with both the bait and prey constructs grew under these stringent conditions. Sequence enumeration of So constructs follow the So-PA protein sequence.

### Molecular cloning of expression vectors and leucine zipper constructs

Standard molecular cloning methods were used to generate recombinant DNA vectors in the pUAST-attB plasmid backbone. Complimentary leucine zipper (ZIP1 and ZIP2; based on those used in split-GAL4 (Pfeiffer *et al*., 2008)), flexible linker (ASG_10_TS) and epitope tag (3XFLAG+3XMYC or 3XFLAG+3XHA) sequences were concatenated and ordered as synthetic gene blocks (IDT and Genewiz/Azenta) to produce pUAST-derivative “ZIP” vectors compatible with N– and C-terminal protein linkages. Vectors containing epitope tags but not ZIPs were similarly constructed. Full length So and Eya cDNA clones were obtained from the *Drosophila* Genomics Resource Center (DGRC) and used to amplify coding sequences with adapters suitable for cloning.

Wobble bases in the N-terminal 129 nucleotide region of the *so* cDNA were edited to synonymous codons to make the sequence resistant to Vienna *Drosophila* Resource Center RNAi reagent 104386 (KK108128); the translated amino acid sequence is unaffected. Synthetic gene blocks containing bespoke or BOS-associated missense mutations were used to modify existing RNAi-resistant vectors.

Though not used in this study, the RNAi-resistance feature is intended to be used in paradigms where So is essential for viability, in future studies. Final vectors underwent plasmid sequencing to verify their integrity; complete vector sequence files and protein translations are provided in Supplemental Information.

### Cell culture

*Drosophila* S2-DRSC (S2) cells (RRID:CVCL_Z992) were purchased from DGRC. Cells were grown in M3 Medium (Sigma) supplemented with 10% heat-inactivated fetal bovine serum (SH30070.02, HyClone), 50 units/mL penicillin G + 50 µg/mL streptomycin sulfate (Gibco) at 25°C. Cell between passages 3-30 were used for experiments. S2 cells were seeded in 6-well (3 x 10^6^ cells in 2.5 mL/well) or 24-well plates (5 x 10^5^ cells in 0.5 mL/well) and were transfected 18 hrs after seeding using TransIT-Insect (MIR 6100, Mirus), according to the manufacturer’s instructions. Plasmids used were pΔTub-GAL4 and pΔTub-hRluc (Zhang *et al*, 2024b), pGL3-*ARE*-Fluc (Silver *et al*., 2003), and pUAST-Eya and / or pUAST-So expression vectors, as indicated in Figures. For luciferase assays, 0.25 ng of pΔTub-hRluc, 50 ng of pΔTub-GAL4, 100 ng of pGL3-*ARE*-Fluc, and 100 ng of the indicated expression vectors were transfected per well in 24-well plates (total 350.25 ng). For co-immunoprecipitation experiments, 200 ng of pΔTub-GAL4 and 750 ng of each expression vector were transfected per well in 6-well plates (total 1700 ng).

### Luciferase assays

S2 cells were lysed at room temperature in passive lysis buffer (Promega) 72 hours after transfection. Firefly and *Renilla* luciferase activities were measured using the Dual Luciferase Reporter Assay System (Promega) using a Synergy 2 plate reader (BioTek). Raw luminescence data were exported and analyzed in Microsoft Excel.

### Drosophila genetics and fly lines

See **Tables S1 and S2** for the sources of stocks used in this study and for precise experimental genotypes represented in figures, respectively. New site-directed transgenic lines were generated commercially (Bestgene Inc.); So-expressing constructs were inserted at ZH-68E (Chr 3) and Eya-expressing constructs were inserted at ZH-22A (Chr 2). Transgene expression was driven in eye tissues by *ey*-GAL4.

In wildtype *Drosophila*, compound eyes are sexually dimorphic in size (Surkova *et al*, 2021). Hence, experimental animals were segregated by sex. Pilot experiments confirmed that genotype effects were equally penetrant in both sexes; thereafter we elected to image only adult female eyes. Animals of both sexes that are homozygous for the *so^1^* allele are viable but eyeless (Cheyette). At most, 30 animals (60 eyes) were imaged from any given biological replicate cross; expected Mendelian ratios resulted in the recovery of fewer progeny in certain experimental crosses. Crosses were repeated a minimum of 2 times.

### Image acquisition and processing

Images of adult flies were acquired on a Nikon SMZ18 microscope using an Accu-scope Excelis-HD camera. Eye area was determined by manual tracing of images in FIJI (NIH) by observers blind to the experimental genotype. Camera images were calibrated using a stage micrometer (ThorLabs). Figures were assembled in Adobe Illustrator.

### Western blots and immunoprecipitation

S2 cells were processed 72 hours post transfection. Prior to protein extraction, some cells were washed in PBS and resuspended in PBS containing 2mM dimethyl 3,3′-dithiopropionimidate dihydrochloride (DTBP), a crosslinking agent (see Figure captions). Reactions proceeded on ice for 50 min before being quenched in 50mM Tris-HCl pH 8.0. Cells were lysed at 4°C in buffer containing 150 mM NaCl, 50 mM Tris, 1% NP-40, 1X cOmplete Mini EDTA-free Protease Inhibitor (04693159001, Roche), 1X Pierce Phosphatase Inhibitor (A32957, Thermo), Phosphatase Inhibitor Cocktail I (10 µL/mL, P2850, Sigma), 5 mM NaF, 1 mM Na_3_VaO_3_, 1 mM β-glycerophosphate, and 1 mM phenylmethylsulfonyl fluoride. Protein extracts were quantified using the Pierce BCA Protein Assay Kit (Thermo). Immunoprecipitation (overnight at 4°C, with rotation) with Pierce α-HA magnetic beads (88836, Thermo) or Pierce α-C-myc magnetic beads (88842, Thermo) was performed with 0.5-1 mg input protein. Sample processing continued as suggested by the bead manufacturer.

Fixed volumes of IP (5-15 µL), or 15-20 µg of input protein were electrophoresed on acrylamide gels (4% stacking, 8% separating) in a Bio-Rad Mini-PROTEAN tank at 100V until the dye front completely transited the separating gel. Protein was transferred to nitrocellulose membrane (1620112, Bio-Rad) using a Bio-Rad semi-dry transfer system for 30-35 min at 15V. Some blots were cut above the 50kDa marker and others above the 25 kDa marker to avoid contaminating signal from IgG heavy and light chains, respectively. For immunodetection of proteins, primary antibodies used were rabbit α-Myc at (71D10, Cell Signaling; 1:4000), rabbit α-HA (C29F4, Cell Signaling; 1:4000) and mouse α-HA (26183, Invitrogen; 1:1000), rabbit α-So (DZ33963, BosterBio; 1:5000) or mouse α-Eya (eya10H6, deposited by N. Bonini and S. Benzer, Developmental Studies Hybridoma Bank, 1:1000). Specific immune complexes were detected with near-IR secondary antibodies (1:10,000), including AlexaFluor680 goat-α-mouse Light Chain (Thermo) and IRDye 680RD or IRDye 800 goat-α-mouse or goat α-rabbit (LiCor). The LiCor Odyssey Imaging System (CLX-0642) was used on auto exposure for both 700 and 800 channels. Images were taken at 169 μm resolution and medium quality. Quantification of signal intensity was performed in LiCor ImageStudio (v5.2).

### Statistical analyses

Luciferase and western blot data were analyzed using one-way ANOVA or one-way normal or lognormal Brown-Forsythe and Welch ANOVA with Dunnett’s T3 correction for multiple comparisons (adjusted *P* values reported), using GraphPad Prism (v11.1.0). The choice of test was driven by the equality or non-equality of within group variance and the normality of the distribution of measurements. Since the bilateral eye tissues of *Drosophila* develop independently of one another, left and right eyes from the same animal were treated as independent observations.

### Rigor and Reproducibility

Representative images and blots are shown in all figures; full blots are shown in Fig. S2. All experiments were repeated at least twice. *Drosophila* genetic crosses were independently executed and scored by multiple investigators. Not all crosses yielded the experimental genotype at expected Mendelian ratios, hence, there are differences in the number of observations between categories. Precise numbers of eyes observed are given in the Figures.

## Results

### Bespoke missense variants protein fragments of So exhibit CoF-selective PPI deficits

The conserved SIX domain of SIX-family TFs is known to mediate PPIs; it consists of a globular tertiary structure comprising 6 α-helices. Using Y2H assays, we determined the site of interaction between Eya/EYA1 and So/SIX1 to reside between the respective EYA domains and SIX domains. More specifically, the interacting portion of the SIX domain was mapped to within the 2 N-terminal α-helices (CP and FP, unpublished data). A co-crystal structure of human SIX1 and EYA2 (PDB: 4egc) supports this model (Patrick *et al*., 2013). In contrast, the N-terminal domain of So was not sufficient to support productive Y2H interactions with either of 2 known repressive CoFs, Groucho (Gro; human TLE1-4) or Sine oculis binding protein (Sobp; human SOBP). Instead, interactions between So and these CoFs required that most of the So or SIX1 SIX domain be present (CP and FP, unpublished data). Alanine-scanning mutagenesis of the human SIX1 SIX domain and analysis of CoF binding profiles by Y2H revealed distinct patterns in the residues affecting EYA1, TLE4 and *Drosophila* Sobp interactions (Neal *et al*., 2024), supporting these unpublished observations.

SIX domain structures and sequences are strongly conserved, with human and *Xenopus* SIX1 SIX domains having identical primary sequences (Pandur & Moody, 2000). Relative to these vertebrate ortholog sequences, the *Drosophila* So SIX domain differs at only 18 of 100 residues (**Fig. S1A**). Focusing on conserved residues within the N-terminal region of the So SIX domain, we engineered a series of missense mutations and performed a Y2H analysis (**Fig. S1B**). What immediately gained our attention were clones So(A109F) and So(C110S) which specifically compromised the So-Gro and So-Eya PPIs, respectively. Henceforth, we will refer to alleles containing these bespoke mutations as SoΔTLE and SoΔEYA, respectively. We selected 3 additional So variants for further analysis – F103E (SoΔEYAΔSOBP), V111E (SoΔPPI) and C112S (SoΔTLEΔSOBP) (**Fig. 1A**). The SoΔPPI allele corresponds to the known human BOS-associated allele SIX1(V17E) (Kochhar *et al*., 2008). Based on their crystallographic data, another group engineered a human SIX1(C16R) allele to specifically disrupt its interaction with EYA2 (Patrick *et al*., 2013). Although we did not test the equivalent *Drosophila* So(C110R) allele by Y2H, we nonetheless included it in select downstream analyses, referring to it as SoΔEYA*.

**Figure 1:**
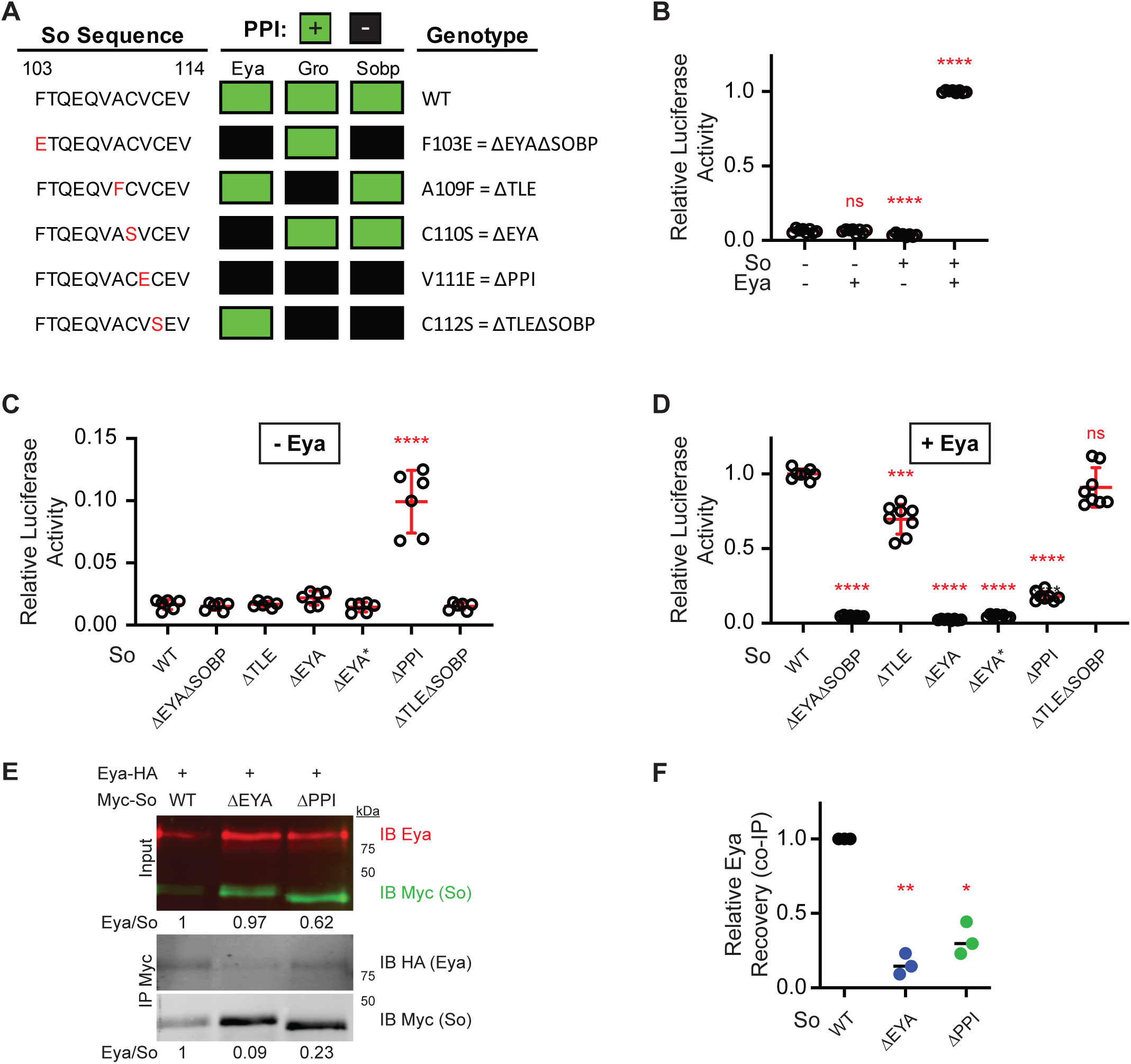
Identification and validation of bespoke PPI-defective variants of So. **(A)** Y2H assays were performed using an N-terminal fragment of the So SIX domain (AA 103-276) and variants (as indicated) with partners Eya (AA 493-766), Gro (AA 1-247) and Sobp (AA 210-749) in pairwise fashion. Positive interactions (green squares) were scored when the interaction supported growth on selective media under conditions in which no background growth was observed for negative controls. Else, no interaction (black box) was recorded. Variant names, ascribed based on So variant PPI profiles, are used throughout the manuscript. **(B-D)** Normalized luciferase activity (p*ARE*-Fluc / p*ΔTub*-hRluc) was recorded from S2 cell lysates; genotypes as indicated. At least 3 biological replicates with 2 technical replicates per trial were assayed and data were analyzed by one-way ANOVA with Dunnett’s T3 posthoc correction (B) or one-way lognormal Brown-Forsythe and Welch ANOVA with Dunnett’s T3 correction (C,D). **** p< 0.0001, *** P<0.001, ns – not significant. The p*ARE*-Fluc reporter responds specifically to the combination of So and Eya, but not either factor alone (B). SoΔPPI exhibited increased basal activity in the absence of Eya, relative to wildtype So and other variants (C); unmarked comparisons in (C) are not statistically significant. In the presence of Eya, variants exhibited different levels of activity (D); those with compromised PPI-Eya had the lowest activity. **(E-F)** Western blot analysis of input and co-immunoprecipitated fractions of S2 cell lysates of the indicated genotypes. Reduced amounts of Eya-HA were recovered as co-immunoprecipitates with SoΔEYA and SoΔPPI, relative to wildtype, in the presence of 2 mM DTBP (E). A representative blot is shown; uncropped blots are provided in Fig. S2. The relative amount of Eya coimmunoprecipitate was calculated from 2 biological replicates of Myc immunoprecipitation and 1 reciprocal HA immunoprecipitation (shown in Fig. S2). Input normalized data are plotted (F) and analyzed by RM one-way ANOVA. ** p= 0.0036; * p=0.0133. The higher electrophoretic mobility of SoΔPPI was consistently observed and may relate to the charge difference imparted by the V111E substitution.

### Bespoke full-length missense variants of So exhibit differences in transcriptional activity and PPIs

Whereas Y2H assays were performed by expressing isolated protein domains or fragments, we wished to evaluate the effects of missense mutations in the context of their full-length proteins. Hence, we cloned wild-type and variant So and wild-type Eya cDNA sequences into standard pUAST expression vectors for reciprocal use in cell culture and in transgenic flies. To ascertain the relative transcriptional activity of So variants, we used a dual-luciferase reporter assay in transiently transfected S2 cells, which do not typically express either So or Eya (https://www.flyrnai.org/cgi-bin/RNAi_expression_levels.pl).

The So+Eya responsive firefly luciferase reporter *ARE*-Fluc was used together with the *Renilla* luciferase control vector, *pΔTub*-hRluc, which is not regulated by So and / or Eya (NJ and SN, data not shown). In control experiments, baseline hRluc-normalized Fluc activity was significantly repressed by the expression of So alone (56.7% of pUAST control, p<0.0001), but not Eya alone (p=0.9977), and was activated greater than 15-fold (p<0.0001) by the co-expression of So and Eya (**Fig. 1B**). Thus, the *ARE*-luc luciferase assay is sensitive to the function of So in the presence and absence of Eya.

Upon expressing So variants alone in S2 cells, SoΔPPI stood apart for having significantly higher basal normalized luciferase activity than the wildtype control (4.5X higher, p<0.0001), or that of any other variant (**Fig. 1C**), although this activity remained well below (∼10%) that induced by wildtype So+Eya (compare scales in **Fig. 1B-C**). Despite being unable to bind EYA2, the corresponding SIX1(V17E) BOS variant was shown to bind a consensus DNA motif with nearly the same affinity as the wildtype SIX1-EYA2 complex using electrophoretic mobility shift assays (Patrick *et al*., 2009). This may account for the higher basal transcription activity that we observed for SoΔPPI.

In the presence of Eya, luciferase results were largely as expected from the Y2H data. All variants with predicted deficits in PPI-Eya (**Fig. 1A**) exhibited severely reduced normalized luciferase activity relative to wildtype So+Eya, including SoΔPPI (17.9% of WT control, p<0.0001) (**Fig. 1D**). For example, SoΔEYA activity was just 2.4% that of WT (p<0.0001). In contrast, SoΔTLEΔSOBP was not expected to have deficits in PPI-Eya based on Y2H evidence and its luciferase activity was not statistically different from wildtype So+Eya (**Fig. 1D**). Somewhat surprisingly, SoΔTLE had 69.0% of the transcriptional activity of the wildtype control (p=0.0009, **Fig. 1D**). This unexpected effect may be the consequence of interfering with the position neighboring the critical Eya-binding residue (Patrick *et al*., 2013).

To determine if the observed luciferase results reflected decreased CoF binding as opposed to the failure of the CoF to successfully transactivate So transcription, we performed co-immunoprecipitation (co-IP) assays; a chemical crosslinker (2mM DTBP) was used to stabilize native protein complexes during immunoprecipitation. For these experiments, So and So variant proteins were appended with N-terminal Myc tags and Eya with a C-terminal HA tag to facilitate immunoprecipitation (collectively “TAG” proteins). This did not confound the previously observed variant effects in luciferase assays (**Fig. S1C**).

A representative western blot of protein lysates from cells co-expressing Eya-HA together with either Myc-So, Myc-SoΔEYA or Myc-SoΔPPI is shown (**Fig. 1E**); full blots are shown in **Fig. S2**. Both So and Eya abundance were increased in variant-containing lysates relative to lysates containing Myc-So. After normalizing the abundance of Eya relative to So in the control input sample, we observed no difference in the ratio of Eya-HA to Myc-So in Myc-SoΔEYA samples (normalized to 1 and 0.97, respectively) and a relative decrease in Eya-HA abundance (normalized to 0.62) in the Myc-SoΔPPI samples (**Fig. 1E**). After immunoprecipitation with α-Myc antibody-conjugated beads, we again normalized the abundance of Eya-HA relative to Myc-So in the control co-IP sample. Myc-So variant yield continued to be enriched relative to Myc-So following co-IP (again normalized to 1). However, we observed a significant decrease in the relative abundance of co-immunoprecipitated Eya-HA with both variants. Across 3 experiments, Myc-SoΔEYA co-immunoprecipitated 15.7% as much Eya as the wildtype control (p=0.0036) and Myc-SoΔPPI co-immunoprecipitated 32.3% as much Eya as the wildtype control (p=0.0133) (**Fig. 1F**). These data show that, in the presence of DTBP, there is some detectable residual binding of Eya to the selected So variants. However, this binding appears to be unproductive with respect to So transcription of the *ARE*-FLuc reporter, as shown above (**Fig. 1D**).

### The So-Eya PPI is necessary for retinogenesis in Drosophila

During retinogenesis in *Drosophila*, So and Eya play a central role in activating several pro-retinal genes. For instance, Atonal expression is induced in the presumptive R8 photoreceptor neuron (Tanaka-Matakatsu & Du, 2008; Zhang *et al*., 2006; Zhou *et al*, 2016; Zhou *et al*, 2014), serving as the trigger for the subsequent development of individual ommatidial clusters of which there are ∼800 in the *Drosophila* wildtype compound eye (**Fig. 2A**). Hence, the formation of ommatidial clusters and the size of the eye itself can potentially be used as proxies for So and Eya function in retinogenesis.

**Figure 2:**
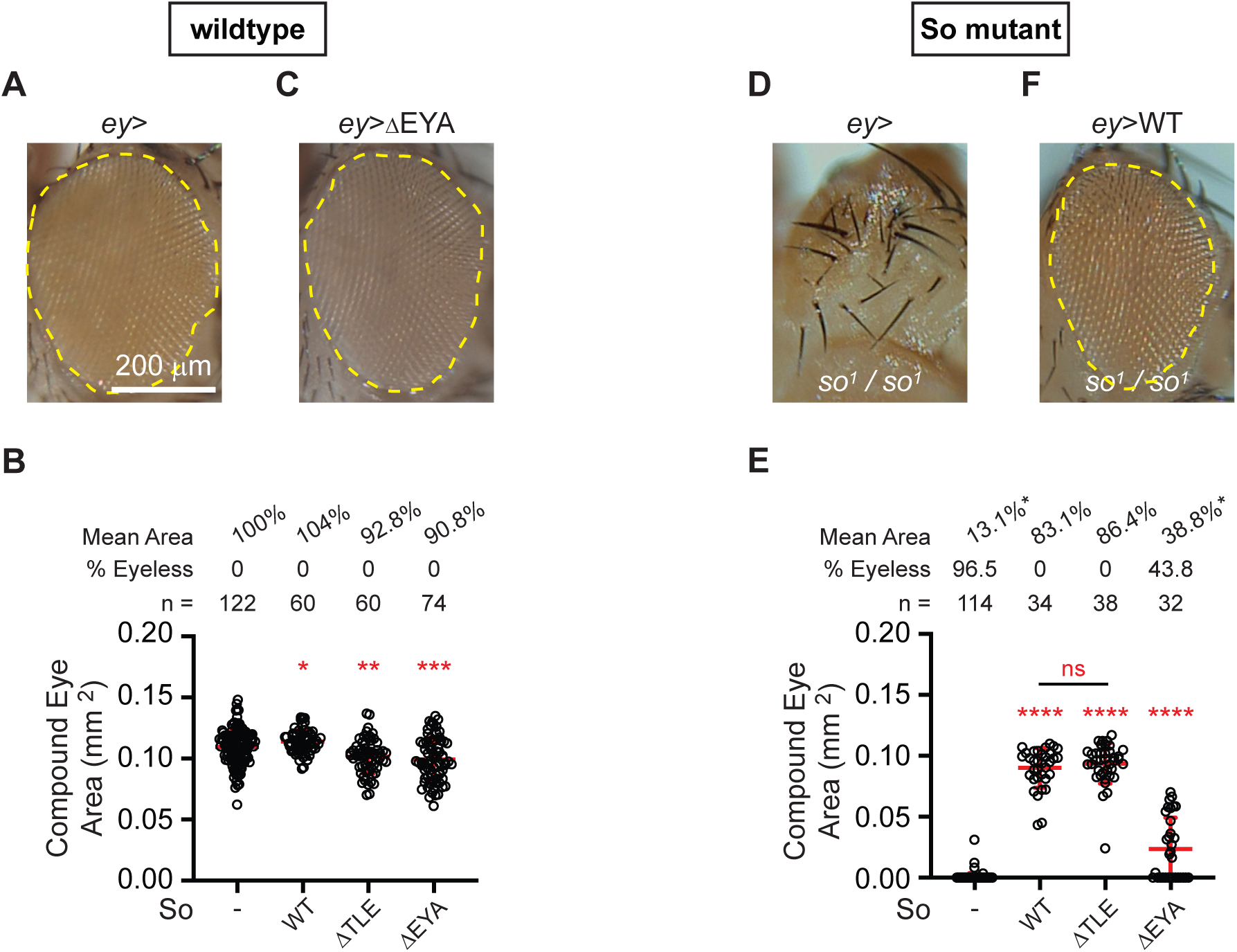
“So mutant” *Drosophila* genetic model for assessing So variant function *in vivo*. **(A)** Compound eye area was quantified from the traced (yellow dashed line) region in a wildtype *Drosophila* compound eye with ∼800 simple eyes, or ommatidia. **(B)** Compound eye area was quantified from flies expressing So variants, as indicated. Data were analyzed by one-way Brown-Forsythe and Welch ANOVA with Dunnett’s T3 correction. *** p<0.001, ** p<0.01, * p<0.05. **(C)** Compound eye from an animal overexpressing SoΔEYA that is smaller than wildtype but not grossly dysmorphic. **(D)** Homozygous *so^1^*/*so^1^*“So Mutant” fly, completely lacking ommatidia. **(E)** Compound eye area was quantified from So Mutant flies expressing So variants, as indicated. Data were analyzed by one-way Brown-Forsythe and Welch ANOVA with Dunnett’s T3 correction. **** p<0.0001, ns – not significant. The percent of eyeless flies is given; mean eye area was calculated only from animals with some eye tissue present. **(F)** Example of a So Mutant fly eye expressing wildtype So, resulting in the restoration of ommatidial development. Scale bar (200 μm) in (A) applies to all images.

To evaluate So and Eya function in eye development, we first generated transgenic flies using the same pUAST vectors as used above in cell culture assays and we used site-directed transgenesis to avert position-effects among allelic variants. We then used *ey*-GAL4 to drive transgene expression in the presumptive eye tissue at a relevant developmental time. In otherwise wildtype flies, expression of SoΔEYA (90.7% of wildtype area; n=74, p=0.0002) or SoΔTLE (92.8% of wildtype area; n=60, p=0.002) each resulted in a slight reduction in eye area while expression of wildtype So resulted in a marginal increase (104.1% of wildtype area; n=60, p=0.0392) in eye area (**Fig. 2B**). The smaller SoΔEYA-expressing eyes do not display gross morphological differences under these expression conditions (**Fig. 2C**). The above results are consistent with an autosomal dominant effect of mutant So, such as those observed for SIX1 variant alleles in BOS patients (Ruf *et al*., 2004).

Flies homozygous for the *so^1^* allele (*so^1^*/s*o^1^*) – henceforth “So mutant” flies – are viable but typically lack eyes altogether (**Fig. 2D**) (Cheyette *et al*., 1994; Serikaku & O’Tousa, 1994). In this experiment, 96.5% of So mutant flies (n=114) were eyeless and the few remaining flies had eyes with a mean area equal to 12.8% that of wt flies (**Fig. 2E**). We adopted this model as a paradigm to evaluate the So-Eya PPI hypothesis. If the So-Eya PPI is indeed necessary for the retinogenic function of So, we would expect So but not SoΔEYA to rescue eye formation. In contrast, there is little evidence for Gro function as a central contributor to retinogenesis in the fly, and thus we would predict that SoΔTLE would rescue eye formation. As a positive control in this assay, we expressed wildtype So and observed that it significantly restored eye formation (n=34, p<0.0001) in all experimental animals (**Fig. 2E-F**). The mean area of rescued fly eyes was 83.1% of wildtype. SoΔTLE also rescued eye formation in 100% of flies (n=38), to a size indistinguishable from those expressing wildtype So (p=0.8286). Consistent with our expectations, 43.8% of SoΔEYA-expressing flies remained eyeless (n=32) and the size of partially restored eyes was just 38.8% of the wildtype rescue size. Statistically, this was a significant rescue of the mutant phenotype (p<0.0001). These observations support the hypothesis that the So-Eya PPI is specifically required for the retinogenic function of So. Moreover, since So-mutant flies naturally express wildtype Eya, it further supports the conclusion that the retinogenic function of Eya is mediated by So, through the So-Eya PPI.

### Leucine zippers can be used to functionally reconstitute the So-Eya transcription complex

Although the above data support our general hypothesis, the experiments above did not directly test whether the So-Eya PPI is necessary for the transcriptional activity of So and / or for its function in retinogenesis. Ultimately, we needed to find a way to restore Eya-binding to So variants that are intrinsically deficient in this ability. To begin, we borrowed from strategies used in synthetic biology to engineer molecular machines and biological processes (e.g. (Plaper *et al*, 2024; Rosa *et al*, 2021; Thompson *et al*, 2012)). The principal strategy we envisaged was to use two heterologous protein domains that interact with one another and to append these to SoΔEYA and Eya, respectively, to generate a heterologous PPI. Leucine zipper domains (ZIPs) have previously been used successfully in fly, to yield the highly successful “split-GAL4” intersectional gene expression toolkit (Pfeiffer *et al*., 2008).

Here, complementary ZIPs were fused to and separately expressed with either the activation domain or the DNA binding domain of the yeast GAL4 transcription factor. Importantly, these studies demonstrated that ZIP expression was well tolerated in *Drosophila* and that it did not induce significant adverse developmental effects therein.

We thus developed our own “PPI-Rescue” approach, starting with a series of pUAST-based expression vectors harboring complementing ZIP sequences (ZIP1 and ZIP2), epitope tags (3XMyc3XFLAG or 3XFLAG3XHA), and a flexible sequence (ASG_10_TS) to which our proteins of interest could be cloned (**Fig. 3A-A’**; see Supplemental Material for sequences). Both N-terminal and C-terminal fusions vectors were generated because, *a priori*, we did not know which configuration would allow our factors of interest to productively interact. To ascertain this, we used our established luciferase assays to test all possible permutations of N– and C-terminally “zipped” proteins and found that the combination of ZIP1-So and EYA-ZIP2 had the highest relative transcriptional activity (NJ and SN, data not shown). Henceforth, we will call these factors simply “ZIP-So” and “Eya-ZIP”.

**Figure 3:**
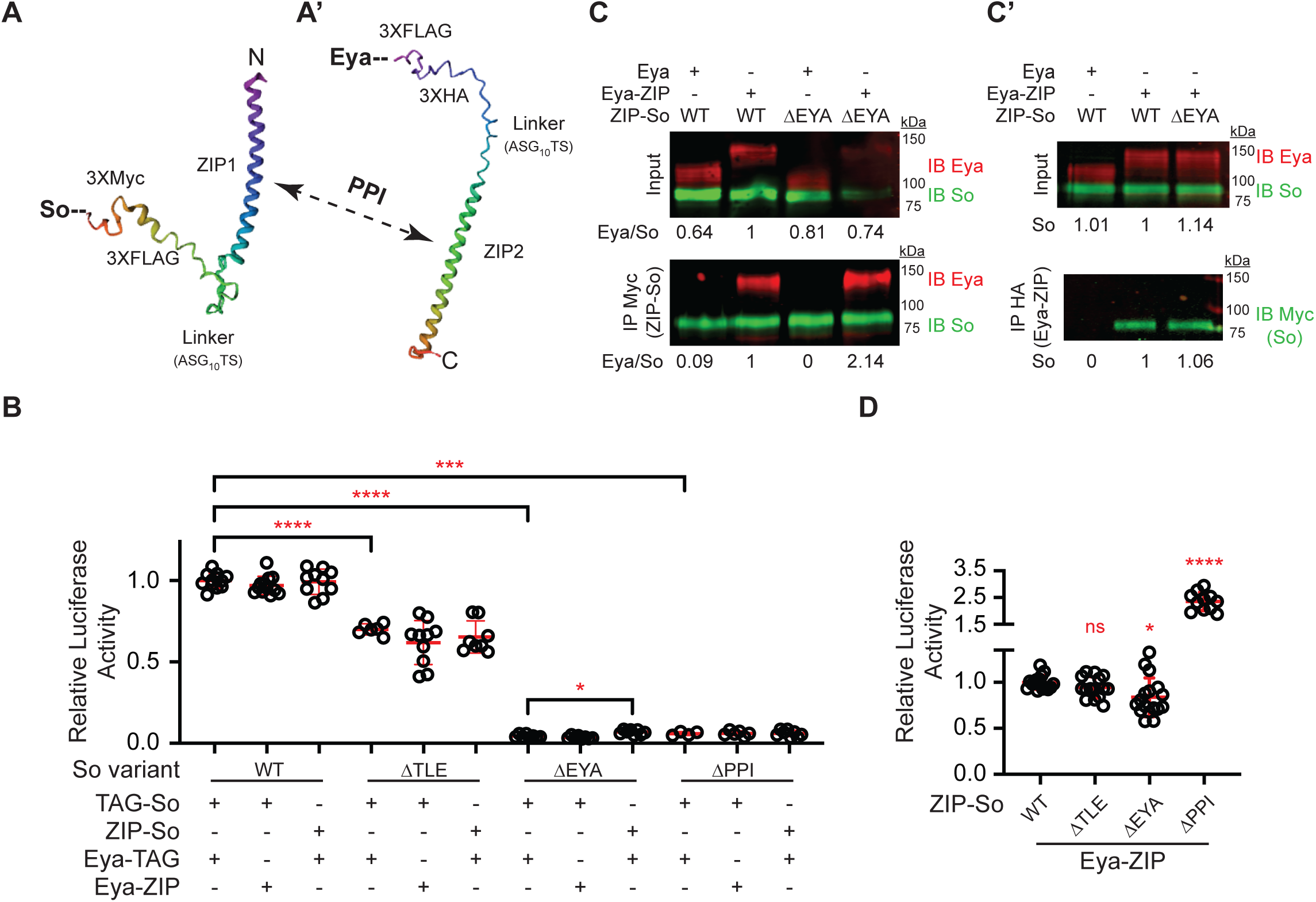
Design and implementation of “PPI-Rescue” to reconstitute a functional So-Eya complex. (A-A’) Leucine zipper domain (ZIP) sequences from split-GAL4 were adapted with epitope tags (as indicated) and used to generate multipurpose cloning vectors. Predicted structures were modeled in RobettaFold (Baek *et al*, 2021). Empirical testing using luciferase assays revealed that N-terminal fusions to So (A) and C-terminal fusions to Eya (A’) functioned in the most complementary fashion. A dashed arrow labeled “PPI” is used to indicate that ZIP1 can dimerize with the complementary ZIP2 domain. **(B)** Luciferase assays were used to demonstrate that ZIP domains do not mask previously observed variant effects on transcription activity. Epitope-tagged (TAG) proteins were used as controls and variant pairs were compared with wildtype to establish the variant effect. Reciprocal pairings of ZIP proteins with TAG partners were then compared to within-variant TAG pairs. One-way Brown-Forsythe and Welch ANOVA with Dunnett’s T3 correction; **** p<0.0001, *** p<0.001, * p<0.05, If not marked, the comparison was not significantly different. **(C-C’)** Reciprocal co-immunoprecipitation assays using ZIP proteins. Myc immunoprecipitations (C) for ZIP-So and HA immunoprecipitations (C’) for Eya-ZIP were performed from S2 cells lysates; genotypes as indicated. In the absence of crosslinker, at least 10-fold more Eya-ZIP than Eya was coimmunoprecipitated with Zip-So (C). Zip-SoΔEYA immunoprecipitated 2.14 times more Eya-ZIP than ZIP-So, compared with a 10-fold reduction with non-ZIP proteins (Fig. 1E). In the reciprocal experiment, Eya-ZIP coimmunoprecipitated equivalent amounts of Zip-So and Zip-SoΔEYA. Note that untagged Eya was used in C’ lane 1 and thus we did not expect any So to be recovered in this sample. **(D)** Luciferase assays were used to demonstrate the PPI-Rescue approach. Wildtype and variant ZIP-So was co-expressed with Eya-ZIP and luciferase activity was recorded. Zip-SoΔTLE activity was not significantly different from wildtype ZIP-So and ZIP-SoΔEYA activity was restored to 83.8% of wildtype. Surprisingly, ZIP-SoΔPPI activity significantly exceeded that of wildtype Zip-So. One-way Brown-Forsythe and Welch ANOVA with Dunnett’s T3 correction **** p<0.0001, * P<0.05, ns – not significant.

To validate the PPI-rescue approach, we initially confirmed that the co-expression of isolated ZIP domains alone or in combination with either ZIP-So or Eya-ZIP did not confer any change in the relevant baseline normalized luciferase activity (NJ and SN, data not shown). ZIPs also did not grossly affect protein stability (see below). Next, we wanted to ensure that ZIPs did not mask previously observed variant effects on luciferase activity. We thus performed combinatorial testing with TAG– and ZIP-fusions of So and Eya. TAGs did not alter variant effects on luciferase activity (**Fig. 3B; Fig. S1C**); Myc-SoΔTLE, Myc-SoΔEYA, and Myc-SoΔPPI all had reduced luciferase activity relative to Myc-So when paired with Eya-HA. When ZIP-So constructs were expressed with Eya-HA, we saw the same expected pattern of luciferase activity (**Fig. 3B**). There was just one statistical difference in luciferase activity, a 55% increase in ZIP-SoΔEYA activity relative to Myc-SoΔEYA when each was paired with Eya-HA (p=0.0111). (**Fig. 3B**). However, this heightened activity remained at just 6.8% of wildtype control levels, consistent with the SoΔEYA activity deficit. When the reciprocal experiment was performed – using Eya-ZIP with Myc-So constructs – no statistical differences in luciferase activity were observed relative to TAG controls (**Fig. 3B**). Thus, ZIP domains do not grossly impair the transcription activity of So or Eya, nor do they mask So variant effects in these assays, when used unilaterally.

We continued our investigation by using pairs of ZIP proteins to determine whether PPI-Rescue could in fact eliminate or reduce variant effects on transcription activity by functionally reconstituting the So-Eya transcription complex. In support of this endeavor, reciprocal co-IPs were used to demonstrate that Zip-So and ZIP-SoΔEya were comparable in their abilities to bind Eya-ZIP (**Fig. 3C-C’**), whereas Eya binding was severely compromised in the Myc-SoΔEYA variant (**Fig. 1E-F**). In the absence of DTBP crosslinking agent, only 9% Eya, relative to Eya-ZIP, was co-immunoprecipitated by ZIP-So (**Fig. 3C**). When ZIP-SoΔEya was used for immunoprecipitation, no measurable Eya was recovered, yet 214% as much Eya-ZIP was recovered, relative to ZIP-So (**Fig. 3C**). In reciprocal co-immunoprecipitations, where Eya-Zip was pulled down, 6% more ZIP-SoΔEya was recovered relative to ZIP-So (**Fig. 3C’**). Thus, PPI– Rescue succeeded in restoring the SoΔEYA-Eya PPI.

We next tested whether PPI-Rescue could restore transcriptional activity to So variant proteins. Excitingly, ZIP-SoΔEYA normalized luciferase activity was restored to 83.8% of wildtype (Zip-So) levels (**Fig. 3D**) in the presence of Eya-ZIP (p=0.0222), whereas Myc-SoΔEYA had just 4% of the activity of Myc-So (**Fig. 1D**). We also observed that ZIP-SoΔTLE normalized luciferase activity was rescued to being indistinguishable from that of Zip-So in the presence of Eya-ZIP (p=0.3441; **Fig. 3D**). These data suggest that Eya recruitment to SoΔTLE and ZIP-SoΔTLE may indeed be slightly compromised. Overall, these data provide strong evidence for the ability of PPI-Rescue to restore transcription complex activity in a targeted fashion.

To address whether PPI-Eya is necessary for So function in this assay, we turned to the ZIP-SoΔPPI variant. As mentioned above, this variant failed to interact with any of the potential partners that were tested using Y2H (**Fig. 1A**). If interactions between So and Gro and / or Sobp are required for transcription activity, Eya PPI-Rescue should fail to restore activity to ZIP-SoΔPPI. Alternatively, if PPI-Eya is sufficient for transcriptional activation in our assay, we should expect to see an increase in relative luciferase activity using Eya PPI-Rescue. To our surprise, Eya PPI-Rescue enhanced ZIP-SoΔPPI normalized luciferase activity to beyond wildtype luciferase activity; it was measured at 236% of ZIP-So levels (p<0.0001; **Fig. 3D**).

Previous reports have shown that EYA fails to localize to the nucleus in cells expressing SIX1(V17E) (Shah *et al*., 2020), corresponding to the SoΔPPI used here. This is one possible mechanism by which decreased transcriptional activity could be observed with this variant. Instead, we observed an exaggerated transcription response when we applied PPI-Rescue to SoΔPPI. While we did not examine Eya nuclear localization here, the fact that we observed transcription implies that So-Eya nuclear function was fulfilled in our assays. If SoΔPPI shares the same heightened DNA-binding affinity of SIX1(V17E) (Patrick *et al*., 2009), the combination of potentiated DNA binding and potentiated Eya recruitment (via PPI Rescue) could explain the increase in target gene expression that we observed. An alternative and not mutually exclusive possibility is that the inability of ZIP-SoΔPPI to bind co-repressors such as Gro and Sobp may also contribute to its higher transcriptional activity. Additional investigations could address these alternatives.

Overall, our data demonstrate that the So-Eya PPI is necessary to maximize So transcription activity in this luciferase assay. Furthermore, they provide validation for the PPI-Rescue approach to study the function of specific transcription complexes.

### The So-Eya PPI is necessary for eye formation in So-mutant flies

Given the proof-of-principle established in our luciferase assays, we proceeded to generate transgenic flies with ZIP-containing expression vectors to further test the PPI-Rescue approach. Using the So-mutant paradigm, the successful application of PPI-Rescue would be marked by our ability to restore eye formation through the co-expression of ZIP-SoΔEYA and Eya-ZIP. However, we first needed to perform numerous controls. For instance, we determined that neither the expression of Eya (n=66, p=0.9822) nor of Eya-ZIP (n=82, p>0.9999) was sufficient to restore eye formation to So Mutant flies* (**Fig. 4A**; \**So Mutant negative control data were pooled across experiments and are also presented in* ***Fig. 2D***). In contrast, the simple expression of wildtype ZIP-So rescued eye formation in 100% of flies (n=77) and yielded an average 89.2% restoration of eye size (p<0.0001) (**Fig. 4A-B**). When ZIP-So was co-expressed with wildtype Eya (n=16), there was no significant difference in eye size relative to ZIP-So alone (p=0.1855) but 12.5% of flies remained eyeless (**Fig. 4A**). In contrast, co-expression of Eya-ZIP with ZIP-So (n=42) significantly increased eye size relative to ZIP-So alone (p<0.0001); resulting eyes were in fact 15.6% larger than wildtype control eyes (**Fig. 4A**).

**Figure 4:**
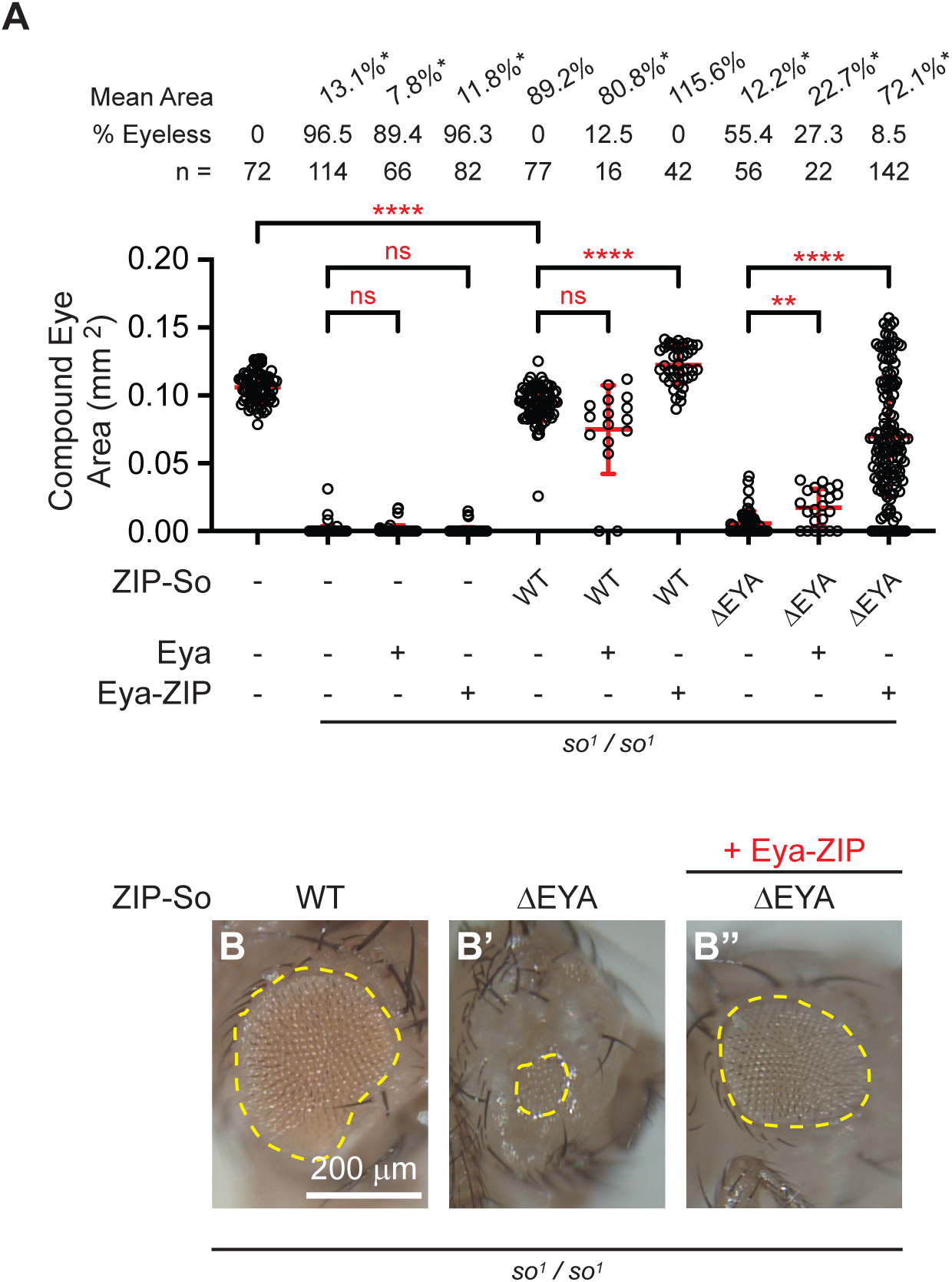
Demonstration of PPI-Rescue in the So Mutant paradigm. **(A)** Compound eye area was measured in flies of the indicated genotypes. In the absence of exogenous So, neither Eya nor Eya-ZIP is sufficient to rescue eye formation in So Mutant flies. Expression of wildtype ZIP-So significantly rescued eye formation in So Mutant flies, but eye size was not completely restored to wildtype levels. When paired with Eya-ZIP, but not Eya, eye size rescue by ZIP-So was improved to beyond wildtype levels. Expression of ZIP-SoΔEYA alone or with Eya only modestly rescued eye development in So Mutant flies. In contrast, co-expression of ZIP-SoΔEYA with Eya-ZIP robustly rescued So Mutant eye development. Mean eye area calculations did not include eyeless flies. One-way Brown-Forsythe and Welch ANOVA with Dunnett’s T3 correction; **** p<0.0001, ** p<0.01, ns – not significant. **(B-B’’)** Representative images of fly eyes from select genotypes reported in (A). Eye area is encircled by a dashed yellow line. Scale bar (200 μm) in (B’) applies to all images.

We next performed the critical series of experiments. We evaluated the ability of ZIP-SoΔEYA to rescue eye formation. When expressed on its own, 55.4% of ZIP-SoΔEYA-expressing flies remained eyeless (**Fig. 4A**), consistent with the inability of SoΔEYA to rescue eye formation (43.8% eyeless; **Fig. 2D**). Rudimentary eyes found in ZIP-SoΔEYA flies were just 12.2% of wildtype size (**Fig. 4A, B’**). The co-expression of wildtype Eya decreased the proportion of eyeless flies to 27.3%, however, the mean size of the rescued eyes was just 22.7% that of wildtype eyes (**Fig. 4A**). Finally, as a true test of PPI-Rescue, we co-expressed ZIP-SoΔEYA with Eya-ZIP. This combination restored eye formation to 91.5% of flies and strikingly, restored mean eye size to be 72.1% that of wildtype (**Fig. 4A, B’’**). We acknowledge that there is a large variance in eye size in PPI-Rescue animals, ranging from the residual 8.5% eyeless flies to the largest observed eye measuring 48.1% larger than the average wildtype eye size.

As for the luciferase data above, our *in vivo* results conclusively demonstrate that the PPI-rescue approach can be used to functionally restore the So-Eya PPI and its related activities, such as activating transcription and promoting eye development.

## Discussion

We have shown that distinct missense mutations in the *Drosophila* TF So can selectively perturb its physical and functional interactions with a limited suite of CoFs. This result has implications with respect to how we should approach the study of BOS (Neal *et al*., 2024), a dominant syndromic disease driven largely by missense mutations in SIX1 and EYA1. Here, by selectively rescuing a specific CoF PPI using heterologous ZIP domains, we have demonstrated that PPI deficits imparted by missense mutations can be physically and functionally restored. The latter is the critical concept of this study – the ability to rescue one specific PPI can serve to demonstrate the necessity and / or sufficiency of a given interaction when it is not feasible to empirically test all possible CoF interactions. Moreover, the paradigm established by having PPI-defective and PPI-Rescue complexes opens the door to molecular genetic, biochemical and omic analyses of transcription complexes and their biological functions.

With respect to So and Eya, our data provides direct evidence for a longstanding hypothesis regarding a role for a So-Eya transcription complex in the *Drosophila* RDGN. We first showed that the SoΔEYA mutation – So(C110S) – eliminated the PPI with Eya in Y2H assays. That the corresponding residue lies at the critical position in the SIX1-EYA2 co-crystal structure reinforces the extreme degree of evolutionary conservation of the SIX and EYA factors (Abdelhak *et al*., 1997; Bellen *et al*, 2019; Bonini *et al*., 1997; Christensen *et al*, 2008; Kawakami *et al*, 2000; Kumar, 2009; Mosrati *et al*, 2011; Neal *et al*., 2024; Yamamoto *et al*, 2014; Zimmerman *et al*, 1997). We next showed that wildtype So, but not SoΔEYA, could activate reporter gene expression in an Eya-dependent manner and that it could restore eye formation in So-mutant flies in the presence of endogenous Eya. To prove that the So-Eya PPI was essential for reporter activation, we used ZIPs to functionally restore transcriptional activity to both SoΔEYA and SoΔPPI, where the latter variant was defective in its interactions with all CoFs that we tested. This evidence demonstrated that a productive So-Eya PPI is necessary for transactivation of So transcription under the prescribed conditions. Finally, we demonstrated PPI-Rescue in transgenic flies where the co-expression of Eya-ZIP with ZIP-SoΔEYA robustly restored eye formation in So Mutants flies.

Ideally, PPI-rescue should not compromise transcription complex activity when both partners are “zipped”. However, we observed ∼70% decrease in ZIP-So + Eya-ZIP normalized luciferase activity, relative to unilateral ZIP controls. One possibility that we considered is that, instead of the desired outcome – to promote a PPI – the presence of both ZIPs may sterically hinder So-Eya complex assembly. Alternatively, the assembled zipper may impede the productive interaction between the So-Eya complex with transcription machinery. Such effects will be important to consider further as we continue to optimize PPI-Rescue. Nonetheless, our application of PPI-Rescue in flies showed that quantitative transcriptional differences observed *in vitro* do not linearly translate to quantitative developmental deficits *in vivo*. That the restoration of eye formation was variable and at times imperfect may also be the result of the broad and constitutive expression of So and Eya driven by *ey*-GAL4. Indeed, many developmental TFs and CoFs are dynamically expressed and thus refinement at the level of expression pattern and / or timing may yield improved consistency in results. Using GAL4 driver lines that are regulated by So enhancers ((Jenett *et al*, 2012)) may help refine PPI-Rescue in the eye, or in other tissues. For example, we noted earlier that Gro and Sobp play only minor roles in retinal development, yet So, Gro and Sobp are known to have important developmental roles outside of the eye (e.g. (Bar-Cohen *et al*, 2023; Birk *et al*, 2010; Chen *et al*, 2008; Choi *et al*, 2005; Jennings & Ish-Horowicz, 2008; Lopez-Rios *et al*, 2003; Tavares *et al*, 2021; Zhu *et al*, 2002)). The reagents and methodology developed here will allow us to investigate the role(s) of So-Gro and So-Sobp transcription complexes in contexts in which their expression naturally overlaps.

The PPI-rescue approach will also be valuable in parsing the complex phenotypes associated with BOS. As described above, BOS manifests in various tissues during development and even individuals carrying the same mutation and within the same family exhibit variance penetrance and expressivity of BOS-associated symptoms (e.g. (Kochhar *et al*., 2008)). While the prevailing hypothesis is that BOS results from the failure of the SIX1-EYA1 transcription complex to function, this is in fact inconsistent with BOS being an autosomal dominant disease. Our previous published data and the data herein demonstrate that a prevalent BOS allele – SIX1(V17E) – has deficits in multiple PPIs (Neal *et al*., 2024).

Others have shown that this variant has higher DNA-binding affinity for its consensus sites and our data provide evidence that the corresponding *Drosophila* variant has heightened basal transcription activity. Since SIX factor DNA-binding is mediated by the homeodomain, located distally to the SIX domain and the specific N-terminal α-helix containing the variant amino acid, it is possible that the lack of PPIs with co-repressors such as TLE1-4 and SOBP account for the observed properties. Using the PPI-rescue approach, this hypothesis could be tested.

Ultimately, we believe that PPI-Rescue reflects a generalizable approach to study the function of discrete transcription complexes in development and disease. We have shown here that it can be used in quantitative functional assays and in a developmental paradigm, and we expect that PPI-Rescue will be equally if not more useful in dissecting complex molecular genetic pathways using omic methods.

## Supplemental Information

**Supplemental File 1.** DNA sequences of plasmid vectors used in this manuscript.

**Supplemental File 2.** Predicted protein translations of the TAG and ZIP proteins.

**Supplemental Table S1.** *Drosophila* stocks used in this study and their sources.

**Supplemental Table S2.** Precise genotypes of animals presented in specific figures.

**Figure S1:**
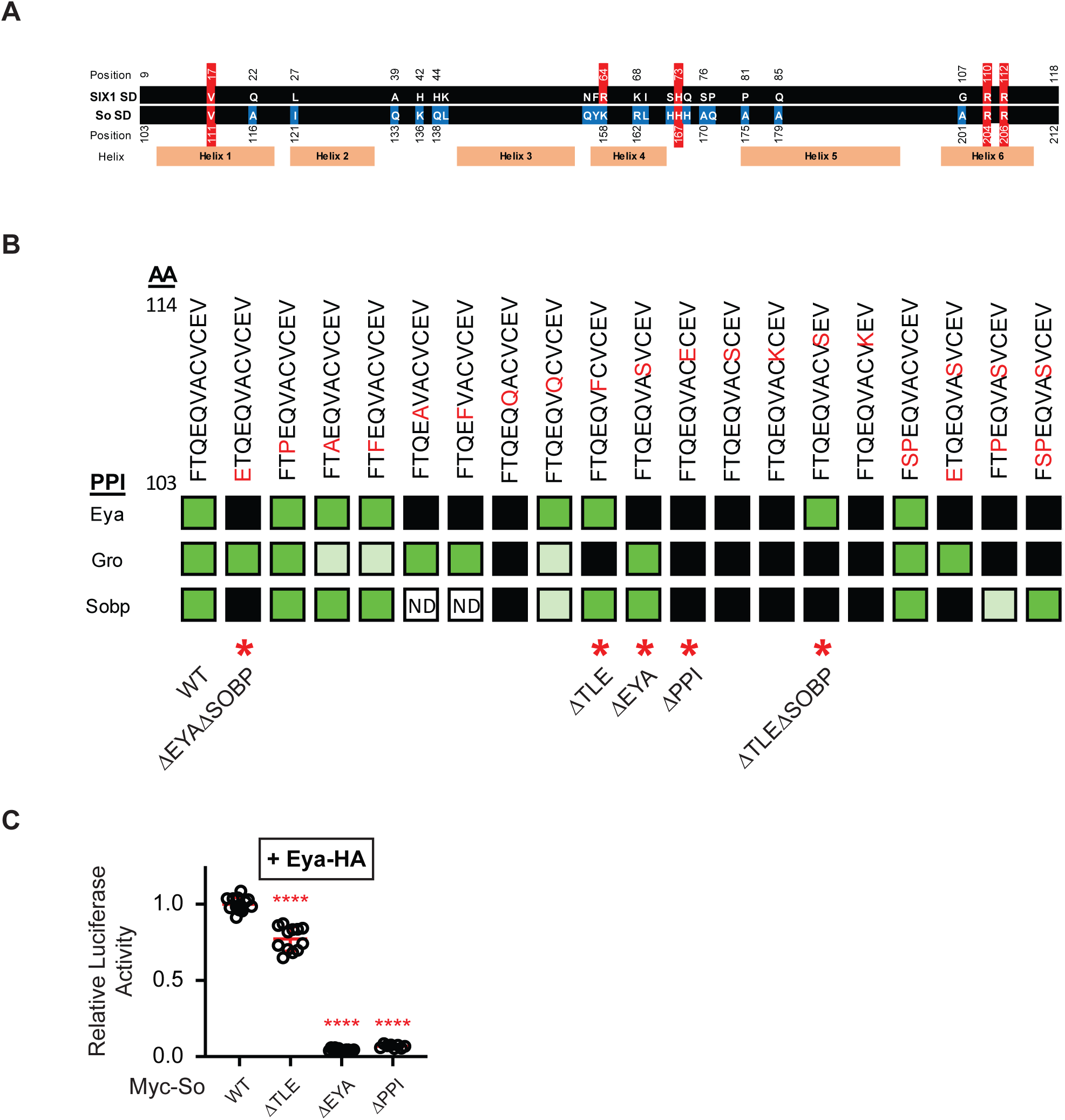
Supplemental data pertaining to Figure 1. **(A)** Schematic comparing the human SIX1 SIX domain sequence with that of *Drosophila* So. Eighteen of 110 residues differ in identity (marked in blue highlight). Residues highlighted in red indicate those associated with BOS (MIM #602588 / 608389 / 113650). The predicted positions of the 6 α-helices are indicated by orange boxes beneath the alignment. **(B)** Extended yeast 2-hybrid data related to Fig. 1A. Variants marked by red asterisk (*) were selected for further study. **(C)** Luciferase assay demonstrating that epitope-tagged Eya and So variants used for immunoprecipitation, exhibit the same activity patterns as their untagged versions. One-way ANOVA with Dunnett’s T3 correction **** p< 0.0001

**Figure S2:**
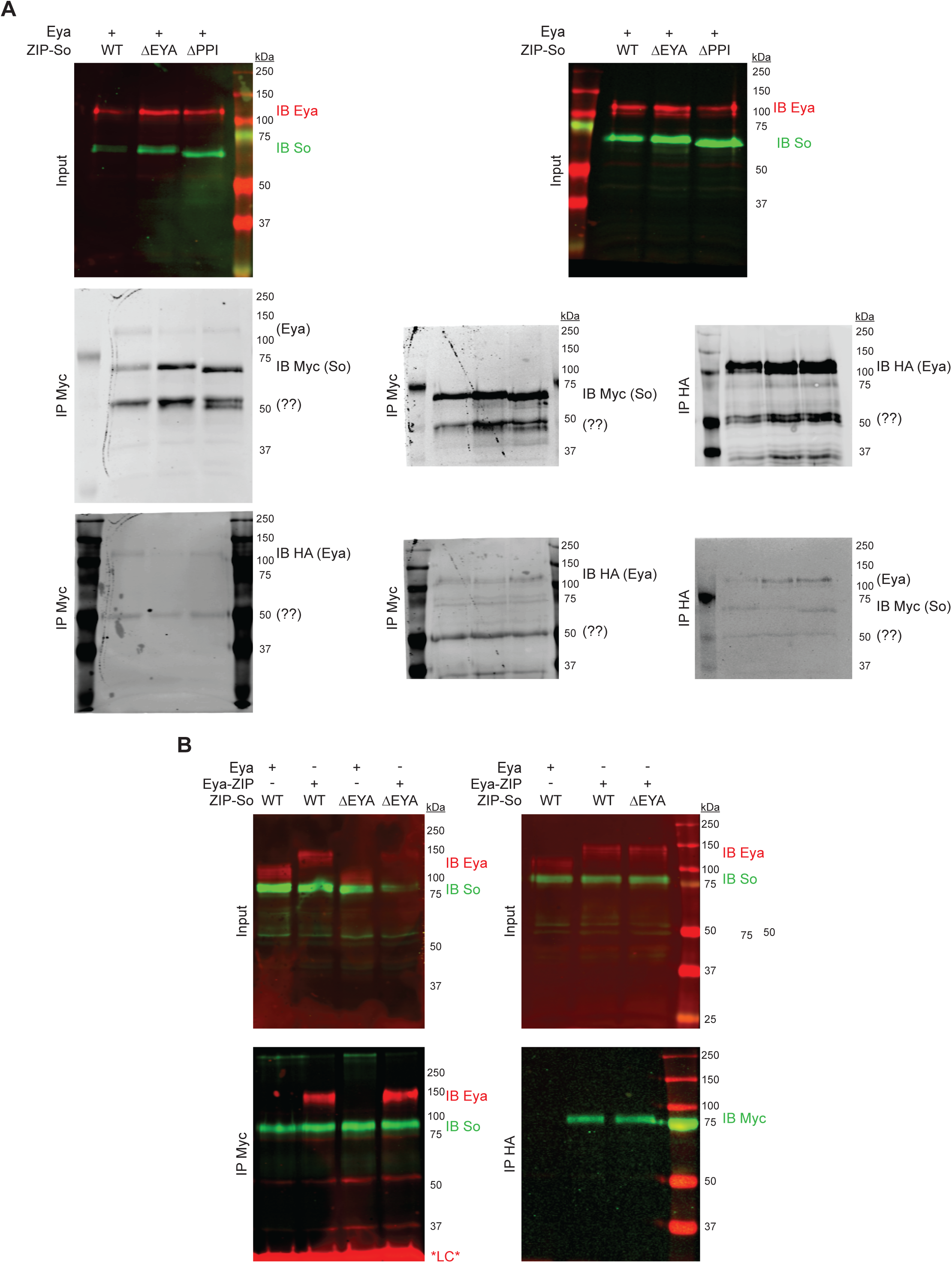
Uncropped western blots. **(A)** Full size blots related to Fig. 1E-F; the representative image shown (Fig. 1E) was derived from the left series of panels and the remainder were used for quantification (Fig. 1F). **(B)** Full size blots related to Fig. 3C (left), and Fig. 3C’ (right).

## Author contributions

Conceptualization: FP, SN

Investigation: NJ, CP, DN, JL, SN

Writing and Editing: SJN

Funding: FP, SJN

## Acknowledgements

This work was supported by grants 5R01EY013167 and 1R03HD094040 (PI Pignoni) and startup funds from Upstate Medical University to SN. The authors wish to thank Christian Diou and Qingxiang Zhou for their effort towards this work.

